# Fine-tuning functional syndromes for stressful environments: lessons on survival from the South African resurrection plant *Myrothamnus flabellifolia*

**DOI:** 10.1101/2021.09.12.459909

**Authors:** Rose A. Marks, Mpho Mbobe, Marilize Greyling, Jennie Pretorius, D. Nicholas McLetchie, Robert VanBuren, Jill M. Farrant

## Abstract

Resilience to abiotic stress is associated with a suite of functional traits related to defense and longevity. Stress tolerant plants are generally slow growing with extended leave lifespans and reduced allocation to reproduction. Resurrection plants are ideal systems to test for trade-offs associated with stress tolerance due to their extreme resiliency. While, growth defense trade-offs are well-characterized, few studies have tested for natural variation associated with tolerating the harshest environments. Here, we surveyed a suite of functional traits related to stress tolerance, leaf economics, and reproductive allocation in natural populations of the South African resurrection plant *Myrothamnus flabellifolia*. We selected three distinct field sites in South Africa ranging from mesic to xeric. Despite considerable environmental variation across the study area, *M. flabellifolia* plants were extremely and similarly stress tolerant at all sites. However, we detected notable variation in other life history and morphological traits. Plants in more mesic sites were larger, faster growing, and had more inflorescences. In contrast, plants from the most xeric sites appeared to invest more in persistence and defense, with lower growth rates and less reproductive allocation. Together, this suggests that desiccation tolerance is a binary trait in *M. flabellifolia* with little natural variation, but that other phenotypes are more labile. The trait syndromes exhibited by plants at the different study sites align with general expectations about growth defense tradeoffs associated with the colonization of extreme environments. We show that plants from the least stressful sites are more reproductive and faster growing, whereas plants from the most stressful sites were slower growing and less reproductive. These findings suggest that *M. flabellifolia* plants are finely tuned to their environment.

## INTRODUCTION

Land plants are incredibly diverse, spanning over 500 million years of evolution and divergence (Kumar *et al*. 2017; Morris *et al*. 2018; Nie *et al*. 2020). As a result, individual species exhibit vastly different life history, anatomy, and physiology from one another. This extensive diversity has allowed plants to colonize a wide array of habitats and coexist in rich communities while simultaneously exploiting unique niches (Whittaker 1965; Huston 1994). Many important adaptations that allow plants to thrive in specific habitats have evolved recurrently and convergently across diverse lineages (Mooney and Dunn 1970; Pérez *et al*. 2004; Lengyel *et al*. 2010; VanBuren *et al*. 2019), and plants with similar and/or complementary traits are often found growing together in close association (Winemiller *et al*. 2015). In fact, the immense variation in plant life strategies can be crudely summarized into three broad functional classes: Competitors, Stress-tolerators, and Ruderals. These classes comprise the basis of CSR theory and can be used to predict species occurrence and distribution (Grime *et al*. 1997; Grime and Mackey 2002; Grime 2006; Pierce *et al*. 2013; Novakovskiy *et al*. 2016; Li and Shipley 2017). The different functional classes exist along a spectrum, but plants in each class are expected to exhibit a characteristic suite of functional traits and dominate under specific environmental conditions. Competitors generally maximize resource acquisition, invest heavily in vegetative growth, and are long-lived. They dominate in low stress, low disturbance sites. Stress-tolerators allocate more resources to tolerance and defense, have slow growth rates, limited plasticity, and are long-lived. They are expected to dominate in high stress, low disturbance sites. Ruderals are short lived, fast growing, and highly reproductive plants that are expected to dominate in low stress, high disturbance sites (Grime *et al*. 1997; Grime and Mackey 2002; Grime 2006; Novakovskiy *et al*. 2016). Few, if any, plants are able to thrive in high stress, high disturbance sites, but in order to do so they would likely employ a combination of both Stress-tolerator and Ruderal functional traits. While the classes defined by CSR theory are a useful tool for predicting functional traits and species distributions (Zhou et al 2017), not all species fall neatly into the classes of Competitor, Stress-tolerator, and Ruderal. Instead, they exist along a spectrum exhibiting complex and variable assemblages of functional traits (Pierce *et al*. 2013). Although the existence of this phenotypic spectrum is widely accepted, few studies have investigated the nuanced variation and functional trade-offs within a single class, or single species (Wellstein *et al*. 2013).

Functional traits and trade-offs associated with stress tolerance are of particular interest due to increasingly challenging environmental conditions related to global change (Ahuja *et al*. 2010). Resurrection plants are, in many ways, the epitome of the Stress-tolerator functional type. They are a phylogenetically diverse group of highly stress tolerant plants that occur in extremely arid habitats across the world (Gaff 1971, 1986; Gaff and Bole 1986; Porembski 2007, 2011; Alcantara *et al*. 2015; Rabarimanarivo and Ramandimbisoa 2019). These plants can tolerate complete desiccation of their vegetative tissues--to or below an absolute water content of -100 MPa--without dying (Bewley 1979). Most resurrection plants can persist in a completely desiccated state for months to years, during which time they may be exposed to intense heat and irradiation, yet they still recover normal metabolic and photosynthetic processes within hours to days of the first rains (Gaff 1989; Farrant *et al*. 1999). Because of their high stress tolerance, resurrection plants are an ideal system in which to test the predictions laid out in CSR theory. Stress-tolerators are expected to dominate in high stress, low disturbance environments, exhibit slow growth rates, minimal reproductive investment, and low plasticity (Novakovskiy *et al*. 2016). However, many resurrection plants are widely distributed and exposed to notable environmental differences across their native range (Marks *et al*. 2021), which could drive intraspecific variability among populations (Stewart and Nilsen 1995; Baythavong 2011). Characterizing the degree of phenotypic variability in resurrection plants will provide insight into the inherent constraints of high stress tolerance and improve predictions on survival of these and other plants.

Here, we tested for phenotypic variation in the resurrection plant, *Myrothamnus flabellifolia* across an environmental gradient. *Myrothamnus flabellifolia* is native to southern Africa and is culturally significant. The plants are highly aromatic and produce a robust profile of secondary compounds related to tolerance and defense (Bentley *et al*. 2020) many of which have important historical and contemporary medicinal applications (Dhillon *et al*. 2014; Jaspal *et al*. 2018; Bentley *et al*. 2019b, 2020). The species is widely known by local people across its native range and has been independently named by multiple communities. It is known as *Uvukakwabafile* in isiZulu, *Patje-ya-tshwene* in Setswana/Sesotho, *Umazifisi* in isiNdebele, and *Mufandichumuka* in Shona. Elders from these communities report using the plant as a medicinal tea to improve the quality of sleep, cure common colds, mitigate fatigue and stress, and as a seasoning for food. The leaves of this plant are also burned, and the smoke is used to treat respiratory ailments such as asthma, to treat headaches and nosebleeds, to revive people who have fainted, and to treat uterine pain. More recently, the plant has been used to treat cancer, reduce blood pressure, and prevent kidney damage (Jaspal *et al*. 2018; Erhabor *et al*. 2020). There is growing international interest in *M. flabellifolia* for cosmetic and pharmaceutical applications, which could lead to over harvest and population decline (Erhabor *et al*. 2020). A better understanding of the ecology of this species is needed to facilitate conservation efforts. Although *M. flabellifolia* plants are generally restricted to sites where abiotic stresses (e.g., aridity, heat, irradiation) are high and disturbance and competition are low, notable environmental variation is evident across its native range (Moore *et al*. 2005, 2007; Bentley *et al*. 2019b). This provides a convenient framework to test for natural variation and trade-offs associated with stress tolerance within a single species. *Myrothamnus flabellifolia* is also dioecious, and sexual dimorphisms may be related to stress tolerance traits, as previously observed in bryophytes (Newton 1972; Stark *et al*. 2005; Stieha *et al*. 2014; Marks *et al*. 2016, 2020). Here, we characterized a suite of functional traits related to stress tolerance, leaf economics, reproductive allocation, and sexual dimorphism in *M. flabellifolia*. We used these data to estimate the degree of phenotypic variability among populations and to test for trade-offs between stress tolerance and other life history traits. We predicted that stress tolerance traits would trade-off with growth and reproduction, as outlined in CSR theory. We also predicted that stress tolerance traits would exhibit the least intraspecific variability compared to growth and reproductive traits due to considerable stabilizing selection on stress tolerance even in the more mesic sites (because drought still occurs at these sites), and that growth and reproductive traits would be more liable.

## MATERIALS and METHODS

### Study organism

*Myrothamnus flabellifolia* is a resurrection plant in the eudicot lineage Gunnerales. Plants are distributed throughout southern Africa in disjunct populations from Namibia to Tanzania, with the highest density of plants occurring in South Africa and Zimbabwe (Fig. 1) (Moore *et al*. 2007; Bentley *et al*. 2019a). *Myrothamnus flabellifolia* is a woody shrub growing up to ∼1.5 M tall with highly branched anatomy, short internode length (∼0.5-1.5 cm), and opposite leaf arrangement. Leaves are small, with short petioles and parallel venation. Inflorescences consist of densely packed florets in a simple arrangement with extremely short pedicles. Male inflorescences have bracts whereas females are bract -less.

**Figure 1.**
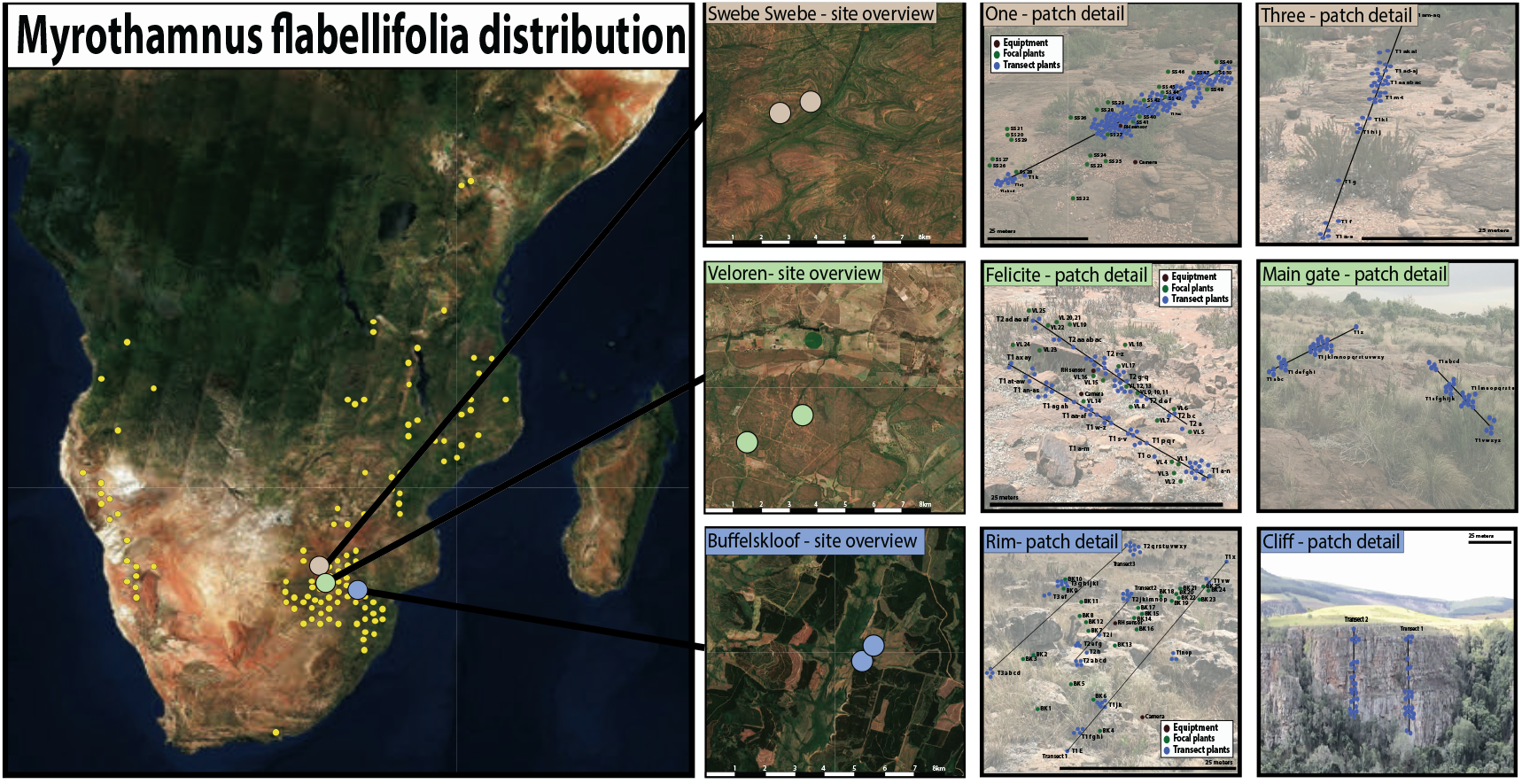
Overview of the study area and sampling design. **a)** The complete distribution of *Myrothamnus flabellifolia* as visualized from data available at www.gbif.org. **b)** The three study sites are located in north eastern South Africa and are within 400 km of one another. **c)** Within each site, two patches of *M. flabellifolia* were sampled, and data on ∼50-150 plants (blue dots) were recorded along transects within each patch. At select patches we tagged 25 focal plants (green dots) for detailed measurements and long term monitoring. Background images of patches are representative photos of each patch and are not scaled to size.

### Study area and environmental characteristics

For the current study, *M. flabellifolia* plants were sampled at three sites in north-eastern South Africa: Buffelskloof Nature Reserve in Mpumalanga (−25.30229, 30.50631), Veloren (−24.7863, 28.3755), and Swebe Swebe Wildlife Estates (−23.7949, 28.0705) in Limpopo (Fig.1). The study sites span 400 km and were selected to capture the maximal environmental variation possible within the study area. Historical weather data from 1981-2019 on temperature, precipitation, and relative humidity was downloaded from NASA’s Prediction Of Worldwide Energy Resources platform (https://power.larc.nasa.gov). Site-specific data was retrieved using the GPS coordinates for the center of each site. These data were analyzed to compute mean annual temperature, rainfall, and relative humidity and to chart monthly temperature and precipitation fluctuations at each of the three study sites. Site-specific elevations were recorded using a Garmin 64csx GPS, bedrock types were visually assessed, and cross referenced with geological records, and soil depths were measured at the base of every study plant (∼50-200 per site). Specimens were vouchered at the Buffelskloof Nature Reserve Herbarium, Lydenberg, Mpumalanga, South Africa (specimen number BNRH0025621).

### Plant sampling

Sampling was done during the rainy season (∼November to March) for two consecutive years (2019-2021). At each of the three sites, two discrete “patches” of *M. flabellifolia* were sampled. We targeted patches that were more than 100 m away from the nearest road (to minimize disturbance) and where *M. flabellifolia* was among the dominant species (based on the criteria that *M. flabefollia* plants must comprise ∼30% or more of the vegetative cover). The location and area of each patch was recorded using a Garmin 64csx GPS. Within each patch, we established transects spanning a cumulative distance of ∼60-100 m. The first transect was set to span the longest axis of the patch and subsequent transects were established until a cumulative distance of 60 m or more was sampled. We sampled all plants that occurred within one meter on either side of each transect for a total of ∼50-150 plants per patch. We targeted plants that appeared to be separate genets with no above ground connections. One of the patches at Buffelskloof is on a vertical cliff, so we sampled plants by rappelling over the cliff and establishing vertical transects following the same procedures described above.

In addition to sampling plants along transects, we selected 25 focal plants at each site for detailed phenotypic measurements. For these, we intentionally targeted plants that were within a medium size range of ∼30-80 cm tall, healthy, and were growing in full sun with minimal evidence of shading or competition. These plants were tagged for long term monitoring and used for measures of growth rate, architecture, specific leaf area (SLA), and stress tolerance phenotyping.

### Stress tolerance traits

We quantified three traits related to water deficit stress and recovery. First, we estimated the intensity of a typical drying event at each site by measuring the relative water content (RWC) of field dry material for the set of 25 focal plants per site. We collected a single terminal twig (∼10 cm long) from each plant in a visually desiccated state. The desiccated leaves were removed from the twig and their mass (fresh mass) was determined immediately after collection using a Frankford Arsenal DS-750 scale. Leaf tissue was then fully submerged in water and placed at 4°C in complete darkness for 48 hours, after which tissues were blotted dry and their turgid mass assessed. Tissues were then transported to the University of Cape Town, dried at 70°C for 2 days and their dry mass was measured. RWC was calculated as [(fresh mass - dry mass)/(turgid mass - dry mass)*100].

Next, we assessed the recovery and rehydration dynamics of desiccated plants. To do so, three randomly selected terminal twigs (∼6-13 cm) were cut from each of the 25 focal plants per site, in a visually desiccated state. Plants were photographed immediately upon collection and placed in individual 50 ml falcon tubes containing 15 ml water each. Plants were imaged again after 2, 4, 8, 12, and 24 hours of rehydration. From these images, we computed the percent of tissue that had recovered for each twig (three replicates per individual, 25 individuals per site, totalling 225 twigs across all three sites). Wez scored each leaf as recovered or not, as this is visually quite distinct, and calculated the percent of recovered leaves relative to the total number of leaves on each twig. Subsequently, we measured the rate of rehydration using the same photographic time series described above. The number of leaves that had rehydrated at each timepoint (2, 4, 8, 12, and 24 hours) was visually determined and used to compute the proportion of leaves open at each time point.

### Growth and leaf traits

Next, we measured a suite of vegetative traits. First, we quantified plant height by measuring the distance from the soil to the top of the tallest branch. Compactness (an anatomical measure of leaf frequency) was estimated for the 25 focal plants per site using photographic data from the rehydration time series described above. Briefly, the number of leaf pairs per cm was determined by counting the number of leaf pairs and dividing that by the total length of the twig using ImageJ v1.53 (Schneider *et al*. 2012). To compute SLA, images of 5-10 fully hydrated leaves per plant were taken and the leaf area was calculated using ImageJ v1.53. The leaf tissue was then dried at 70°C for 2 days and the dry mass assessed. From these measures, we computed SLA as (leaf area / dry mass). Growth rate was estimated for the 25 focal plants per site using three randomly selected terminal twigs per plant. In year one, we marked each twig 10 cm from the tip. The following year, we measured the distance from that mark to the tip of the twig. Any distance beyond 10 cm was considered new growth. Unfortunately, we were only able to gather growth rate data for two of the three sites, since extensive floods in 2020 prevented access to Swebe Swebe during the critical sampling period.

### Reproductive traits

Male and female *M. flabellifolia* plants have distinct floral morphology and thus sex was determined visually when plants were in flower. Population sex ratios were calculated as number of males relative to the total number of reproductive individuals. To investigate differences in reproductive allocation across sites we estimated the number of inflorescences produced by each plant (n=525) at the beginning of the rainy season. Because *M. flabellifolia* can produce dozens of inflorescences on a single plant, we subsampled plants by randomly selecting three apical branches on each plant and counting the number of inflorescences on the top 10 cm of each branch.

### Statistical analyses

Initially, we tested for differences in environmental conditions (e.g., rainfall, temperature, relative humidity, and soil depth) across sites using mixed effects linear models. Next, we tested the effect of site, sex and their interaction on relative water content using a mixed effects linear model. Percent recovery data were log transformed to improve normality and the effects of site, sex, and their interaction were tested using a mixed effect linear model. To test for differences in the rate of rehydration, we used MANOVA repeated measures analysis to test the effect of site and sex on the proportion of leaves open over time. For plant height, compactness, and SLA we used mixed effects linear models to test the effects of site and sex on each response variable, respectively. For growth rate, we used student’s T-test to identify significant differences between the two sites where data were available. A heterogeneity test was used to determine if population sex ratios were different from one another and a goodness of fit test was used to identify sex ratios significantly different from 50:50. Lastly, we ran a mixed effects linear model to test the effect of site, sex, and their interaction on the number of inflorescences produced. For all models, plant ID and twig replicate were included as random effects.

To gain insight into trait relationships, trade-offs, and associations, we conducted multivariate analyses on all vegetative and reproductive traits. We generated covariance and correlation matrices for each pairwise combination of traits to identify the most positively and negatively associated traits. We also conducted principal component analysis (PCA) to visualize sample and trait relationships. These analyses were conducted using the set of 25 focal plants per site and include trait values for height, compactness, leaf area, SLA, growth rate, and flower production. We did not include stress tolerance traits in these analyses because they showed minimal differences across the study sites.

## RESULTS

### Environmental differences across sites

Substantial environmental differences were detected across the study sites in South Africa (Fig. 2a). Annual rainfall differed significantly (F_2,113_=52.1, P<0.0001), ranging from 820±41.5 mm at Buffelskloof (the wettest site), to 507±26.3 mm at Veloren (the intermediate site) to 430±23.1 mm at Swebe Swebe (the driest site). Significant differences in mean annual relative humidity (F_2,113_=204.2, P<0.0001) and temperature (F_2,113_=289.7, P<0.0001) were also detected along a similar gradient. Relative humidity ranged from 61±0.7% at Buffelskloof, to 47±0.6% at Veloren and 45±0.6% at Swebe Swebe. Temperatures ranged from 18±0.1°C at Buffelskloof, 20±0.1°C at Veloren, and 21±0.1°C at Swebe Swebe. These sites also occur at different elevations and on different bedrock types. The various substrates can contribute to differences in hydrology, mineral nutrient availability, and thermal dynamics, which further differentiate the ecological conditions across these sites. Buffelskloof is situated on quartzite at 1,492 M, Veloren on sandstone at 1,334 M, and Swebe Swebe on conglomerate sandstone at 1,068 M. Soil depth was low across all sites, with mean depth ranging from 6.04±0.35 cm at Buffelskloof, to 7.5±0.38 cm at Veloren and 6.13±0.21 cm at Swebe Swebe, but differences across sites were significant (F_2,210_=6.79, P=0.0014). Somewhat counterintuitively, the cliff population at Buffelskloof had the deepest soil depth, but this consisted of sporadic pockets of deep soil separated by vertical rock. Taken together, these variables combine to generate perceptibly different environmental conditions at the three study sites (Fig. 2b). Buffelskloof is the wettest, coolest, and highest elevation site; Veloren is intermediate on all three measures, and Swebe Swebe is the driest, hottest, and lowest elevation site.

**Figure 2.**
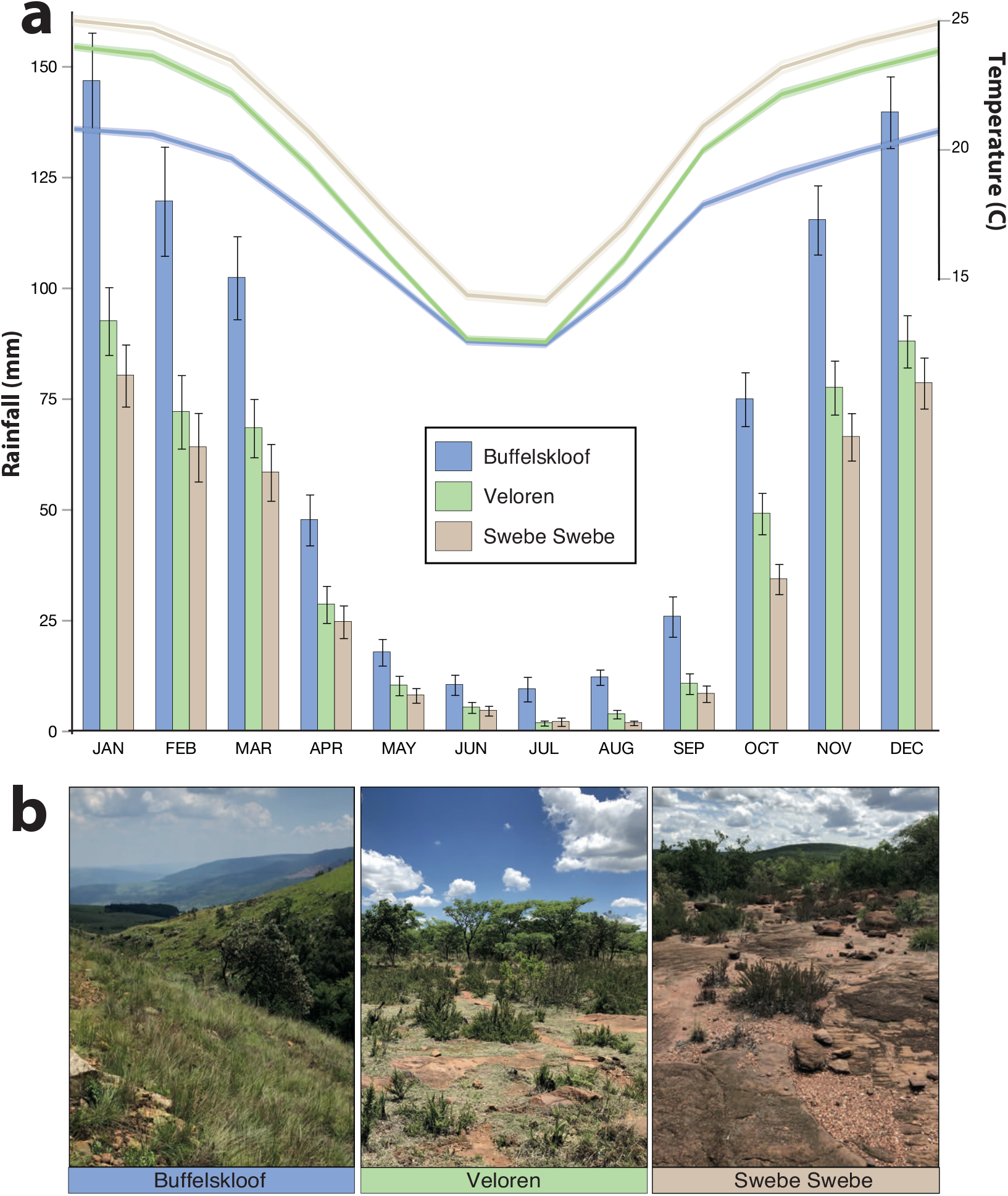
Environmental conditions at the three study sites. **a)** Differences in mean monthly temperature (lines) and rainfall (bars) are evident across the three study sites. **b)** All three sites are classified as dry winter climates, but differences in vegetative cover and community are evident.

### Stress tolerance traits

Desiccation tolerance traits exhibited minimal variation in *M. flabellifolia* plants across the three study sites. Field desiccated plants had a mean RWC of 9.6±0.9% at Buffelskloof, 9.0±0.8% at Veloren, and 9.8±0.9% at Swebe Swebe (Fig. 3a). An exceptionally high percentage of tissue recovered for all plants at all sites with mean percent recovery of 99.95±0.03% at Buffelskloof, 99.53±0.16% at Veloren and 99.89±0.05% at Swebe Swebe and there were no significant differences across these sites (Fig. 3c). The rate of rehydration (measured as the proportion of leaves open at a given time point) was the most variable stress tolerance trait we measured, and despite a small effect size showed significant differences across sites (F_2,192_=2.35, P=0.0021). This appears to be driven by differences at early timepoints where *M. flabellifolia* plants from Veloren were the slowest to rehydrate and plants from Swebe Swebe were the fastest (Fig. 3d).

**Figure 3.**
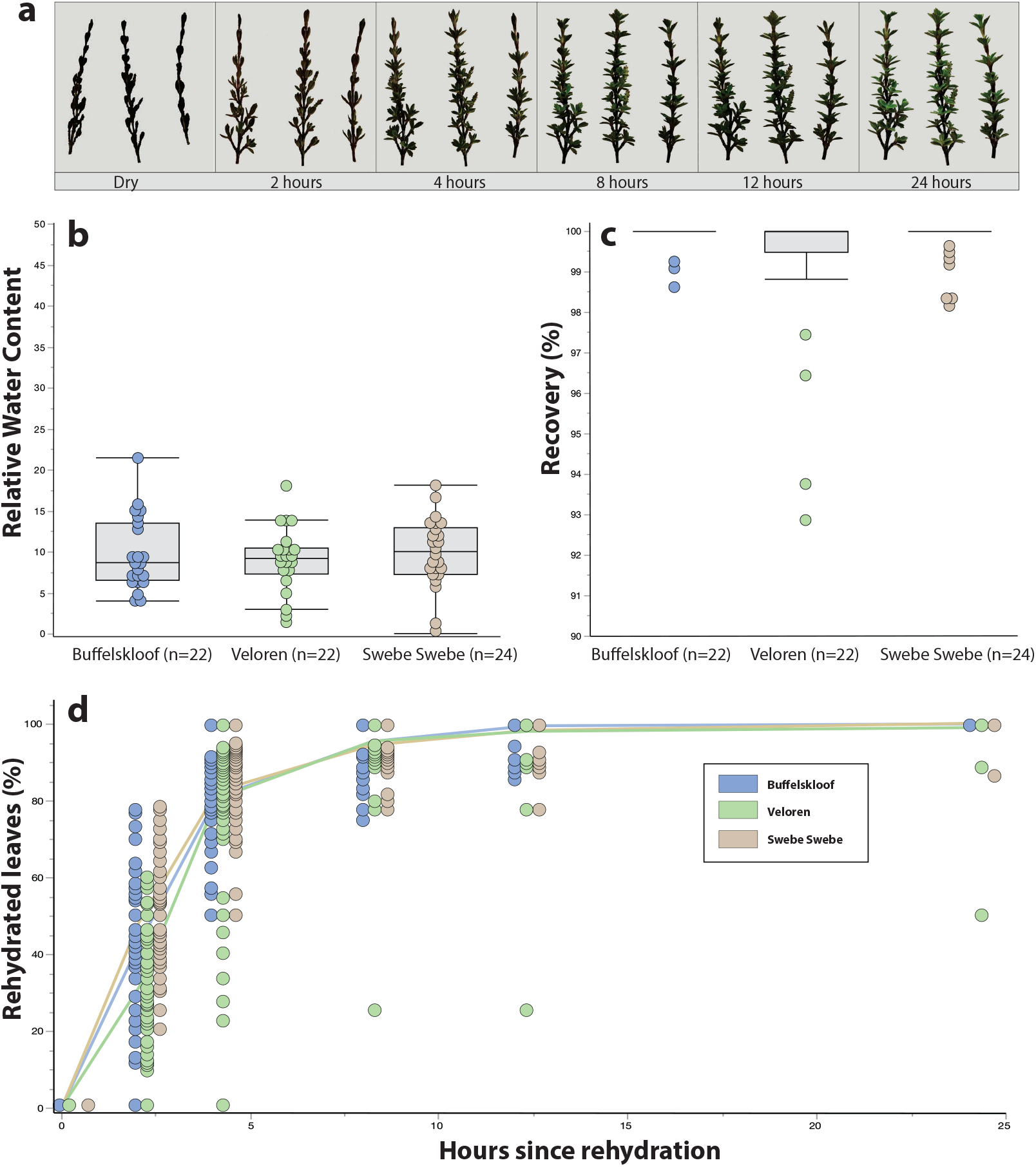
Stress tolerance traits across the three study sites, ordered from wettest to driest. **a)** Pictorial representation of a rehydration time course of twigs from one representative plant from Buffelskloof. These, and similar images for all plants, were used to estimate the proportion of leaves open at each timepoint and to assess percent recovery. **b)** The relative water contents of plants collected from the field in a desiccated state. There were no significant differences across the sites. **c)** The percentage of leaves that recovered to a green and healthy condition within 24 hours. There were no significant differences across the sites. **d)** The rate of rehydration is plotted as the proportion of leaves open (green and healthy) at discrete time points following rehydration. Plants from Veloren were significantly slower to rehydrate than plants from Swebe Swebe and Buffelskloof.

### Growth and vegetative traits

Vegetative traits exhibited more variability than stress tolerance traits (Fig. 4). Significant differences in plant height (F_2,193_=6.11, P=0.0025), compactness (F_2,193_=6.11, P<0.0001), growth rate (t_144_=5.35, P<0.0001), and SLA (F_2,63_=3.24, P=0.046) were identified across sites. Plants at Buffelskloof were the shortest (35.3±2.15 cm), those at Veloren were intermediate (45.9±2.30 cm), and plants at Swebe Swebe were the tallest (55.7±1.68 cm). Plants at Veloren had the most compact leaf arrangement (1.4 ±0.04 leaves per cm) but the lowest SLA (8.9±0.39), whereas those at Swebe Swebe were least compact (1.09±0.03 leaves per cm) but had the highest SLA (9.43±0.38) and plants from Buffelskloof were intermediate for both (1.28±0.04 leaves per cm and 8.80±0.26 SLA). Plants from Buffelskloof grew significantly more in one year (5.80±0.39 cm) than plants Veloren (2.17±0.33 cm).

**Figure 4.**
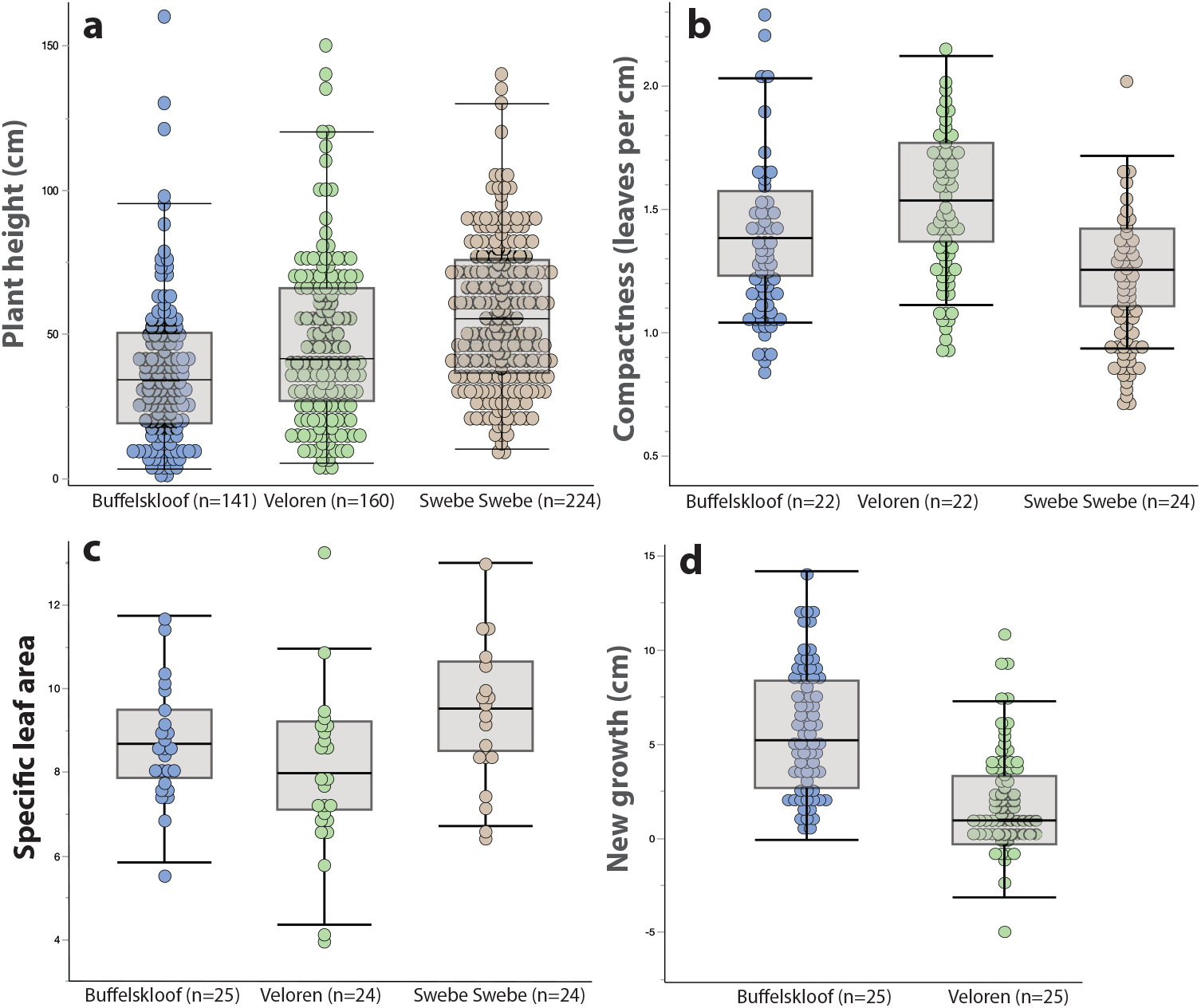
Vegetative traits across the three study sites, ordered from wettest to driest. **a)** Plant height differed significantly across populations. **b)** The number of leaves per cm (a measure of compactness) differed significantly across the study sites. **c)** Specific leaf area differed significantly across sites. **d)** The amount of annual branch growth (a proxy for growth rate) differed significantly between Buffelskloof and Veloren. We were unable to measure growth rate at Swebe Swebe because extensive flooding in 2020 prevented access to field sites during the critical sampling window.

### Reproductive traits

We identified significant differences in the number of inflorescences produced at each site (F_2,447_=22.8, P<0.001). Plants from Buffelskloof produced the most inflorescences (3.8±0.29 inflorescences per 10 cm), plants from Veloren were intermediate (3.2±0.22 inflorescences per 10 cm), and plants from Swebe Swebe produced the fewest (1.7±0.12 inflorescences per 10 cm).

### Trait associations and tradeoffs

Multivariate analyses were used to identify trait correlations and covariances. Principal component analysis (PCA) was conducted to visualize sample relationships and gain insight into trait syndromes and functional classes. We identified several traits with strong positive and negative associations (Fig. 5). The most positively associated traits were growth and inflorescence production (fast growing plants produced more inflorescences) and height and leaf area (taller plants had larger leaves). Our analyses identified multiple negatively associated traits, which point towards possible trade-offs. Height and inflorescence number (taller plants produced fewer inflorescences), height and SLA (taller plants had lower SLA), and SLA and inflorescence number (plants with higher SLA produce fewer inflorescences) were all negatively associated, suggesting that being tall and having low SLA trades-off with reproductive allocation. PCA reinforced these associations showing that growth and inflorescence number project in the same direction and are opposed to SLA, height, and compactness (Fig. 5). Plants from Swebe Swebe formed the most distinct cluster and are characterized by increased height and SLA, with fewer inflorescences and open leaf arrangement. Plants from Buffelskloof are the most reproductive and fastest growing plants.

**Fig. 5.**
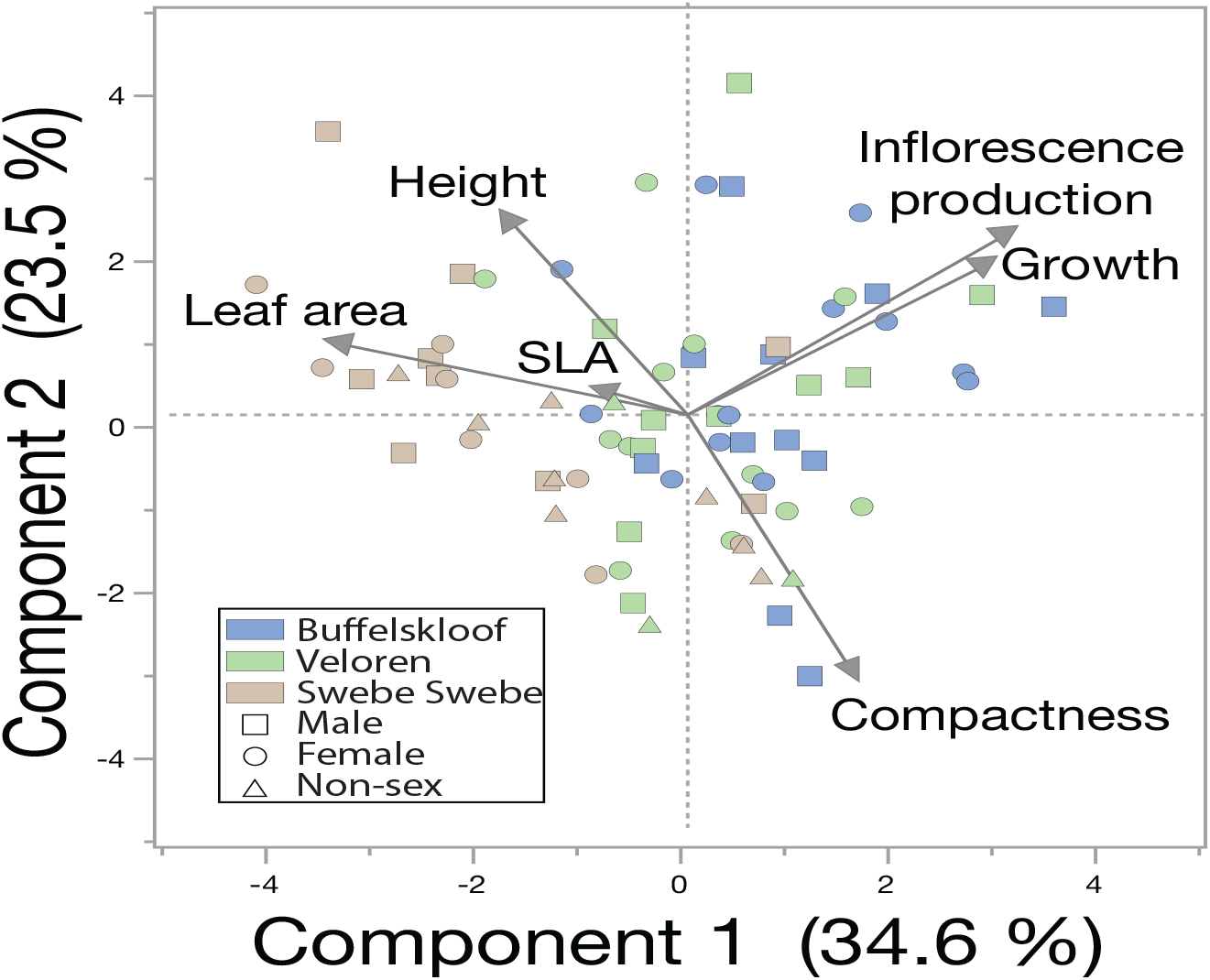
Principal component analysis of six traits for the 25 focal plants per site. Trait projections are plotted on the graph as arrows and each plant is indicated by a single point. Points are colored by study site and shaped by sex.

### Sexual dimorphisms

Sampling was done during the rainy season when flowering is most common. Consequently, we were able to sex a relatively high proportion of plants (60-70% in most sites) (Fig. 6a). The only exception were cliff dwelling plants at Buffelskloof, which had much lower flowering rate, possibly due to limited light exposure. We detected variable sex ratios across patches and sites. Buffelskloof and Veloren were slightly (but not significantly) female biased, whereas Swebe Swebe was significantly male biased (G_1_=4.19, P=0.04) (Fig. 6b).

**Figure 6.**
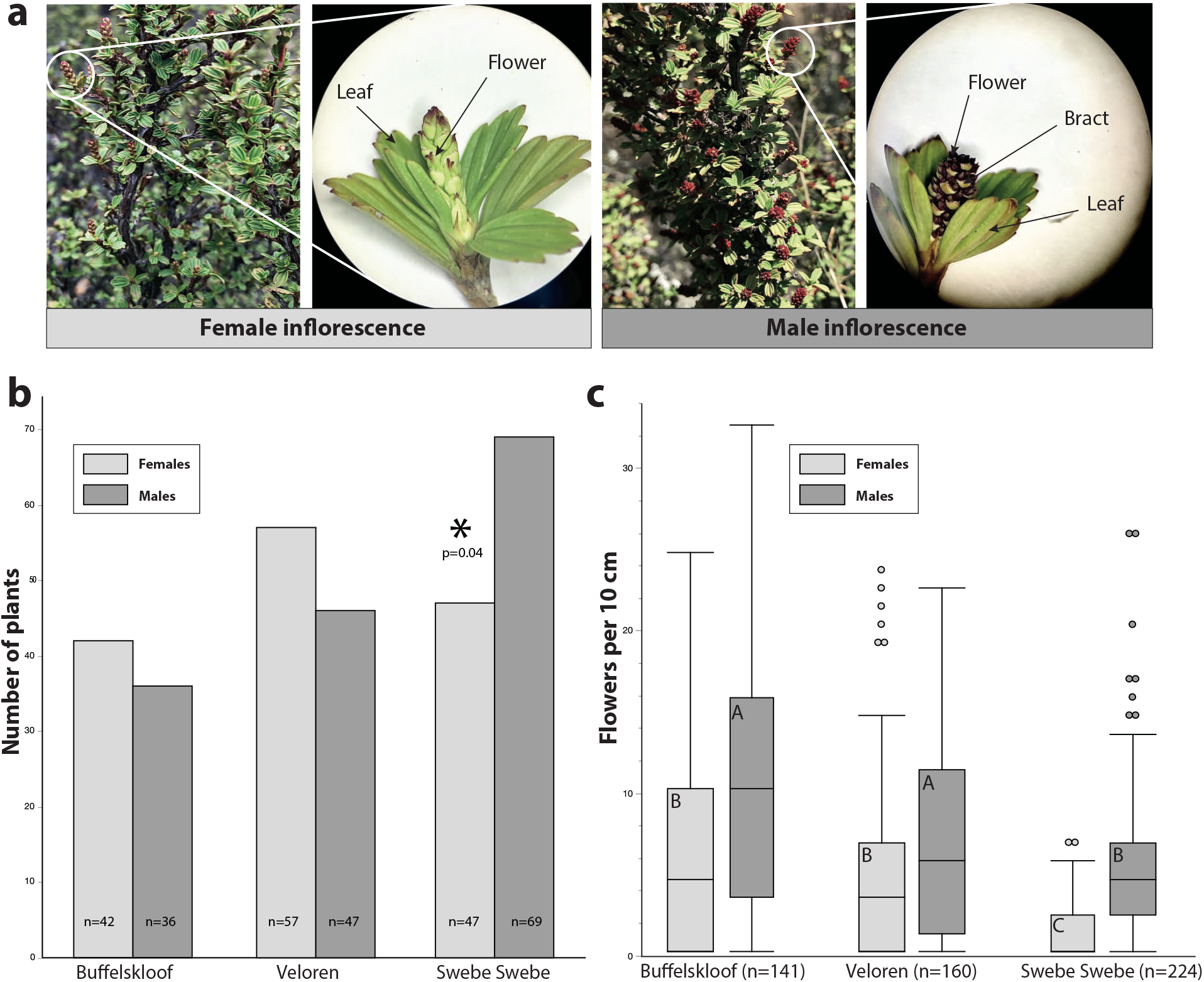
Reproductive traits and sex ratios across the three study sites, ordered from wettest to driest. **a)** Floral anatomy of female inflorescences (left panels) and male inflorescences (right panels). **b)** The number of male and female plants at each site (non-sex plants are not shown). Swebe Swebe was significantly male biased. **c)** The number of inflorescences on male and female plants. Males produced more inflorescences than females at every site. Different letters in box plots show significant differences among sites, corrected for multiple comparisons by Tukey’s HSD.

We did not detect sexual dimorphisms in any stress tolerance or vegetative traits. Male and female plants had equivalent field RWC, rehydrated at the same rate, and had similarly high percentages of recovery. There were no significant differences between the sexes in height, compactness, SLA, or growth rate. However, we did detect a sexual dimorphism in inflorescence production. Males produced significantly more inflorescences than females across all sites (F_2,603_=211.6, P<0.0001) (Fig. 6c).

## DISCUSSION

In theory, stress tolerance is associated with a suite of functional traits related to defense and longevity, including slow growth, extended leaf-lifespan, reduced allocation to reproduction, and minimal plasticity (Grime 2006; Stanton *et al*. 2000). *Myrothamnus flabellifolia* exhibits many of these characteristic phenotypes, including low growth rates (plants put on only 3.34±0.37 cm of annual branch growth compared to elongation rates of 5-15 cm reported for other species (Rossatto *et al*. 2009; Rossatto 2009) and low SLA (mean SLA is 8.3±0.29 in *M. flabellifolia* relative to 25.6±1 0.63 reported in a meta analyses across functional classes (Pierce *et al*. 2013)). In addition, *M. flabellifolia* plants were found to be extremely stress tolerant across all study sites. Despite marked environmental variation across the study area, we detected astonishingly little variability in tolerance, suggesting strong stabilizing selection on desiccation tolerance even in more mesic areas.

Despite *M. flabellifolia’s* classic Stress-tolerator phenotype, we detected complex and noteworthy phenotypic variation in vegetative and reproductive traits across the study area. Plants from the wettest study site (Buffelskloof) were small, semi-compact, fast growth, and highly reproductive, giving them a somewhat Ruderal-like strategy. Plants from the intermediate site (Veloren) were tall, extremely compact, had slow growth rates, and intermediate flowering-- a more typical Stress-tolerator syndrome. Plants from the driest site (Swebe Swebe) were tall and open, with high SLA and minimal flowering--exhibiting an almost Competitor-like strategy. Taken together these differences point towards the existence of subtle sub-syndromes within a single Stress-tolerator species. Because stress tolerance exhibited so little variation across the study sites, we cannot conclusively identify trade-offs with any other traits. However, the different sub-syndromes exhibited by plants at the three sites hint at a possible stress-tolerance productivity trade-off. Plants from the most mesic (presumably least stressful) of the study sites, exhibit the highest growth rate and potential reproductive output, whereas plants from the more xeric sites had lower reproductive output and reduced growth rates. The fine-tuning of these syndromes to specific habitats has major implications for species survival and adaptation to changing environments (Negreiros *et al*. 2014)

The Buffelskloof syndrome is the most Ruderal of the three sites, with small, small-leaved plants exhibiting high investment in growth and reproduction. The fast growth rates but small size of plants at Buffelskloof suggest that these plants experience considerable turnover or mortality during establishment and growth. Although we did not quantify it directly, competitive interactions are likely a bigger factor at Buffelskloof compared to the other study sites, since vegetative cover is much higher at Buffelskloof (Fig. 2b). This could explain the abundance of relatively small (presumably young) plants and perhaps even the high investment in sex--as a potential escape strategy. Alternatively, the small size of these plants could be impacted by “desiccation pruning” a phenomenon in which plants self-prune because they are unable to refill their xylem above a certain height.

In comparison, plants from Veloren exhibit a more characteristic Stress-tolerator phenotype; they grow slower, have low SLA, and invest less in reproduction. This integrates nicely into CSR theory by showing that in increasingly dry conditions, *M. flabellifolia* exhibits a characteristic Stress-tolerator syndrome. Plants from Veloren were also some of the slowest to rehydrate, possibly driven by mechanical dynamics in which taller and more compact plants rehydrate slower due to the higher volume of vegetative tissue that needs refilling.

At Swebe Swebe, the driest of the three sites, the story is a bit more anomalous. These plants actually exhibit the most Competitor-like syndrome of all three sites. In fact, PCA analyses indicate that the plants from Swebe Swebe are the most distinct, despite the environment being somewhat similar to Veloren. At Swebe, plants are tall, have large leaves, and reduced allocation to reproduction. It is unexpected to observe this syndrome in the most putatively stressful site, but it suggests that the Swebe Swebe plants may have a different approach to dealing with sporadic water availability. Perhaps the more open architecture and higher SLA at Swebe Swebe allow plants to cool more effectively in high temperatures and rehydrate more rapidly than those at other sites (Fig. 3d). This strategy might maximize photosynthetic productivity in an area with the most sporadic water availability.

Sex ratio variation is influenced by numerous factors, including stochastic dynamics. In most angiosperms, females are expected to be less stress tolerant due to increased resource demand for reproductive processes (Barrett and Hough 2013; Juvany and Munné-Bosch 2015). Thus, females are expected to be less abundant in stressful sites. However, the extent to which this prediction is upheld by empirical data is highly variable and species-specific (Marks *et al*. 2019). Here, we find that the driest site, Swebe Swebe, is male biased, fitting the expectation. Sexually dimorphic traits, especially those related to stress tolerance, can differentially impact the survival of male and female plants and drive biased population sex ratios and spatial segregation of the sexes, both of which have consequences for population dynamics (Bierzychudek and Eckhart 1988; Juvany and Munné-Bosch 2015). Identifying such dimorphisms is an important step to understanding how to conserve and protect dioecious plants (Petry *et al*. 2016). Sexual dimorphisms are expected to be most obvious in sex related traits (Barrett and Hough 2013), and we detected a single sexual dimorphism in a reproductive trait, but not in any vegetative or stress tolerance traits. We found that males produced more inflorescences than females at all sites. This finding is not surprising, as males typically invest more in pre-fertilization reproductive processes, whereas females tend to incur major costs post-fertilization (Bateman 1948).

In conclusion we show that stress tolerance phenotypes are extremely conserved in *M. flabellifolia*. Despite considerable environmental variation, plants are extremely and equally stress tolerant at all sites. However, we detect notable variation in other life history and morphological traits, suggesting that these plants are finely tuned to their environment. In fact, the trait syndromes exhibited by plants at the different study sites generally align with our expectations about stress tolerance and productivity trade-offs. We show that plants from the least stressful sites are more reproductive and faster growing, whereas plants from the more stressful sites are slower growing and less reproductive. Taken together, these data simultaneously support and deviate from classic CSR theory. We find evidence of classic Stress-tolerator phenotypes (slow growth, low SLA) but we also detect variability and plasticity in other traits, indicating that these plants, despite being extremely stress tolerant, can adjust to different environmental conditions.

## AUTHOR CONTRIBUTIONS

RAM, JMF, RV, and DNM conceived of the study. RAM, MG, and JP collected field data. RAM led data analyses and writing of the manuscript. MM collected and summarized ethnobotanical information. All authors contributed to data interpretation and edited the manuscript.

## ACKNOWLEDGEMENTS

This work was supported by an NSF Postdoctoral Research Fellowship in Biology to RAM (PRFB-1906094) and by the Plant Resiliency Institute at Michigan State University. JMF acknowledges funding supplied by the South African Department of Science and Innovation and National Research Foundation (Grant No. 98406). This work would not have been possible without the generous support of landowners who provided accommodation and valuable local expertise. We thank Syd Carlton and Felicite Jackson for allowing access to Veloren Estate, Ken Maude, Peter and Nadine Vervoot, for access to Swebe Swebe. We thank Barbra Turpin, John, and Sandy Burrows at Buffelskloof Nature Preserve for invaluable assistance in taxonomic identification. Technical and logistical assistance was provided by Keren Cooper.

## Notes

### Competing Interest Statement

The authors have declared no competing interest.

